# Horizontal Gene Transfer Inference: Gene presence-absence outperforms gene trees

**DOI:** 10.1101/2024.12.27.630302

**Authors:** Swastik Mishra, Martin J. Lercher

## Abstract

Horizontal gene transfer (HGT) is a fundamental driver of prokaryotic evolution, facilitating the acquisition of novel traits and adaptation to new environments. Despite its importance, methods for inferring HGT are rarely systematically compared, leaving a gap in our understanding of their relative strengths and limitations. Validating HGT inference methods is challenging due to the absence of a genomic fossil record that could confirm historical transfer events. Without an empirical gold standard, new inference methods are typically validated using simulated data; however, these simulations may not accurately capture biological complexity and often embed the same assumptions used in the inference methods themselves. Here, we leverage the tendency of HGT events to involve multiple neighboring genes to assess the accuracy of diverse HGT inference methods. We show that methods analyzing gene family presence/absence patterns across species trees consistently outperform approaches based on gene tree-species tree reconciliation. Our findings challenge the prevailing assumption that explicit phylogenetic reconciliation methods are superior to simpler implicit methods. By providing a comprehensive benchmark, we offer practical recommendations for selecting appropriate methods and indicate avenues for future methodological advancements.

## Introduction

Horizontal Gene Transfer (HGT) is recognized as a major driving force in the evolution of prokaryotes. (Arnold et al., 2021) However, the inference of HGT events is challenging, since HGT is commonly followed by other evolutionary processes such as sequence amelioration and gene loss. In the best-case scenario, HGT events can be traced between closely related genomes in lab evolution experiments to validate HGT inferences. This limits any rigorous validation of HGT inference methods to only extremely recent HGT events. However, on longer timescales, transfers occur frequently even between distantly related organisms (Cordero and Hogeweg, 2009, Sheinman et al., 2021), something that laboratory evolution experiments cannot replicate. Although one can test the robustness of the HGT inference methods, for example by data resampling, that does not validate the model choices made during the inference process (Felsenstein, 2003).

As a result of these challenges, HGT inference methods are typically validated using simulated data. Simulated data offers key advantages over empirical data for method validation: it provides a known ground truth and allows direct assessment of robustness and sensitivity by varying parameter values or violating specific assumptions. However, simulations are based on a set of assumptions about the evolutionary process, potentially biasing their results in comparison to real data – even if these assumptions are accepted principles or “natural interpretations” of the community (Feyerabend, 1993). Examples of potentially biasing assumptions are: considering genes as evolutionary units, ignoring intragenic rearrangements; neglecting population-level processes such as drift; assuming that sites evolve independently from each other and from structural constraints; and ignoring extinct or unsampled species. Using simulations to compare different HGT inference methods is complicated by the differences in assumptions made by each method: a particular method will perform well in the simulated world designed for it. The vast majority of methods, when published, have not been tested on a common reference set of simulated data, nor have they been compared to one another. Moreover, methods not tested in the real world may fail to capture real-world complexities (Kapust et al., 2018). It is thus valuable to validate and compare methods based on their performance on the same empirical data set.

In the following, we develop a comparative analysis of HGT inference methods using genomics data. Our goal is to provide an approximate ranking of HGT inference methods, delineating which methods infer results that are meaningful in a given biological context, as well as to have a nuanced view of how these methods compare to each other.

HGT inference methods can be broadly classified into two categories: *parametric* (or sequence composition) methods and *phylogenetic* (or tree-based) methods (Ravenhall et al., 2015).

Parametric methods operate under the assumption that the *sequence composition* of the transferred gene differs between the donor and recipient organisms. Sequence features of a genome, such as GC content, codon usage, and oligonucleotide frequencies, are often analyzed. However, they are limited in their effectiveness when the donor and recipient are closely related and thus have similar sequence features, or when dealing with ancient HGT events where enough time has passed for the acquired nucleotide sequence to evolve to the sequence features of the host genome (sequence amelioration).

Phylogenetic methods, in contrast, analyze sets of sequences by leveraging phylogenetic trees. These methods can be categorized into explicit and implicit approaches.

Explicit phylogenetic methods compare the branching patterns (topologies) of gene trees, which show the history of individual genes, with species trees, which show how species are related based on vertical inheritance. When the topology of a gene tree deviates from that of the species tree, such discrepancies can not only indicate the occurrence of an HGT event, but also help infer the direction of transfer – identifying both the likely donor and recipient species. For example, imagine a gene tree where a gene from species A is very similar to genes from a distant clade that contains species B. This mismatch suggests that species A likely received the gene from the lineage of species B through HGT. In the simplest case of a single HGT event, the method reconciles the gene tree with the species tree by introducing a horizontal branch connecting the donor and recipient lineages. However, in bacterial evolution we often need to consider multiple events of duplication, transfer, or loss (DTL) to fully explain the observed differences between gene and species trees, and there may be several plausible reconciliation scenarios for any given pair of trees. The frequency with which a particular DTL event appears across these scenarios reflects the confidence of the method in that inferred event. For example, a transfer event that appears in 90% of the reconciliations indicates a high confidence in that event. Among explicit methods, tools such as RANGER-DTL (Bansal et al., 2018) and ALE (Szöllosi et al., 2012, Szöllősi et al., 2013) use a maximum likelihood (ML) framework to estimate the rates of duplication, transfer, and loss events that best explain the observed gene and species trees. These methods optimize DTL rate parameters to maximize the overall likelihood, and then sample reconciliation scenarios probabilistically according to these rates. In contrast, methods like AnGST (David and Alm, 2011) operate within a maximum parsimony (MP) framework, inferring DTL events by minimizing a cost function for each type of event.

Implicit methods avoid such a direct comparison and hence do not require a gene tree. Instead, they aim to infer the gene gain and loss events that explain the observed gene distribution across genomes in a genome tree. BLAST-hit methods such as DLIGHT (Dessimoz et al., 2008) find very similar gene sequences in distantly related organisms. Methods based on phyletic patterns such as GLOOME (Cohen et al., 2010) and Count (Csűös, 2010) use a statistical framework instead.

Since implicit phylogenetic methods do not require a gene tree, they are faster to execute and are not compromised by the potential inaccuracies of gene tree reconstruction. However, unlike explicit methods, they can only infer the *gain* or *loss* of a gene in a given recipient branch of the species tree, without providing information about the donor of the gene. Additionally, one can save time by not estimating a species tree beforehand and instead let for example GLOOME infer the species tree topology based on the phyletic pattern itself, even if this may result in a less accurate tree topology. Note that all phylogenetic inference methods assume the existence (and correctness) of a species tree describing purely vertical inheritance; however, such a species tree may not always be well defined, especially in cases of highly reticulate evolution.

For a more comprehensive review of HGT inference methods, we refer the reader to Ravenhall et al. (2015).

Although phylogenetic methods offer valuable insights, they are not without limitations. Unrecognized paralogy resulting from duplication followed by loss can be misinterpreted as HGT. Furthermore, tree reconstruction errors, particularly in gene trees, can lead to false HGT inferences (Than et al., 2007). However, with increases in computing power, studies on HGT have increasingly used phylogenetic methods, not least because they are not compromised by sequence amelioration and can thus infer older HGT events than parametric methods.

Parametric methods were developed first and have been benchmarked elsewhere (Becq et al., 2010), with oligonucleotide signature-based methods found to outperform alternative approaches. We thus use an oligonucleotide signature-based method, Wn (Tsirigos and Rigoutsos, 2005), as a representative parametric method for our benchmark, and focus primarily on the comparison of phylogenetic methods. Explicit phylogenetic methods included in our study are ALE, RANGER-DTL, and AnGST. For RANGER-DTL, in addition to the full version, we also include a heuristic ‘fast’ version that samples a smaller space of optimal reconciliations. Implicit methods included in our study are GLOOME and Count, two phyletic pattern-based methods. GLOOME and Count were each run with both ML and MP settings. With GLOOME, each of those were also run with and without an input species tree. Unless stated otherwise, we refer to GLOOME (ML or MP) as versions with an input species tree. Our selection of HGT inference methods for the benchmark aims to provide a comprehensive overview of different state-of-the-art methods. We preferentially include methods that are widely used by the community and have user-friendly, publicly available implementations. There are many other methods that have not been included in this study. These include methods that have no publicly available implementations of the algorithm (e.g., DLIGHT (Dessimoz et al., 2008)); methods that cannot handle modern large datasets with multiple genes of the same gene family in a taxon (especially subtree-prune-and-regraft based methods, e.g. RIATA-HGT (Nakhleh et al., 2005), SPRIT (Hill et al., 2010), EEEP (Beiko and Hamilton, 2006)); methods that focus on the presence of HGT in the evolution of a gene but not on the inference of exact transfer events (e.g. JPrime (Sjöstrand et al., 2014); and additional methods discussed in (Poptsova, 2009)).

Here, we benchmark the selected methods by evaluating how well they reproduce expected patterns in empirical data. Specifically, we expect that genes transferred together in a single HGT event often arrive on the same DNA fragment and initially remain neighbors in the recipient genome. Over time, genomic rearrangements may separate them, but a substantial fraction should still be found close together in extant genomes. Thus, if we find a higher percentage of neighboring gene pairs among the genes inferred by a given method to have been acquired around the same time (i.e., on the same branch of the species tree), this method likely captures true HGT events more reliably than alternative methods. This principle forms the core of our comparative analysis.

Although intuitively, the usage of explicit gene tree information may seem to be an advantage, our findings indicate that implicit phylogenetic methods infer more biologically meaningful HGT events than explicit phylogenetic methods or parametric methods, having a lower false positive rate.

## Results

For our benchmark of HGT inference methods, we used the dataset of orthologous gene families in Gammaproteobacteria in the EggNOG database v6 (Hernández-Plaza et al., 2023). EggNOG terms these gene families non-supervised orthologous groups, or *NOGs*. We focussed on the 359 Gammaproteobacterial taxa in EggNOG that are featured in the tree of life provided by ASTRAL (Zhu et al., 2019); we used the subtree with these taxa as the input genome tree for the phylogenetic methods. We then sampled a representative set of 1286 gene families (NOGs) that occur in at least 10 of these taxa. We obtained the corresponding gene trees from EggNOG, rooting them using Minimum Ancestor Deviation (Tria et al., 2017), which has been shown to be more accurate than other rooting methods (Wade et al., 2020). We used these gene families to infer HGT events using each of the tested methods.

For each method, one can restrict the set of HGT inferences to the most reliable ones, either by modifying an input parameter (the gain/loss penalty ratio for Count (MP) and GLOOME (MP)) or by removing inferences labeled by the inference method as less reliable (e.g., based on the frequency of gain events across reconciliation models by ALE). In this way, each method’s number of HGT inferences can be varied, with lower numbers corresponding to higher “stringency”, i.e., more reliable inferences. Because each method quantifies confidence differently, these thresholds are not directly comparable across methods. To enable fair comparisons, we instead use the numbers of inferred co-acquired gene pairs as common reference points at which we compare the methods. For the explicit phylogenetic methods, we defined stringency as the mean number of transfers inferred per reconciliation model. In GLOOME (ML) and Count (ML), we used the minimum posterior probability of gene gain events as the stringency. For GLOOME (MP) and Count (MP), we use the gain/loss penalty ratio as the stringency parameter. In the parametric method Wn, we fixed the oligonucleotide window size to 8 and defined stringency as the ratio of absolute deviation to the median absolute deviation (MAD; see Methods for details).

### Implicit and explicit methods infer very different sets of HGT events

To assess similarities and differences in the sets of HGT events inferred by each method, we first clustered the methods based on the similarity between pairs of these sets. As a similarity measure, we used the Overlap Coefficient, which is the ratio of the size of the intersection to the size of the smaller set in the pair. Because for each method the set of infered HGT events can expand or contract massively by changing the stringency, we measured similarities as the maximum possible Overlap Coefficient across all possible pairs of thresholds.

As expected, the explicit phylogenetic methods (ALE, RANGER, AnGST) cluster together, as do the implicit phylogenetic methods (GLOOME, Count) (Fig. 1). The biggest differences in terms of inferred HGT events are seen between explicit and implicit phylogenetic methods. The parametric method Wn clusters with the implicit phylogenetic methods. Note that the maximum Overlap Coefficient between all the implicit methods is 1 (or very close to 1 when involving GLOOME ML and MP without a species tree), i.e. for each pair of methods, there are thresholds such that one set of HGT inferences is a subset of the other. The same is true for ALE and AnGST.

**Fig. 1.**
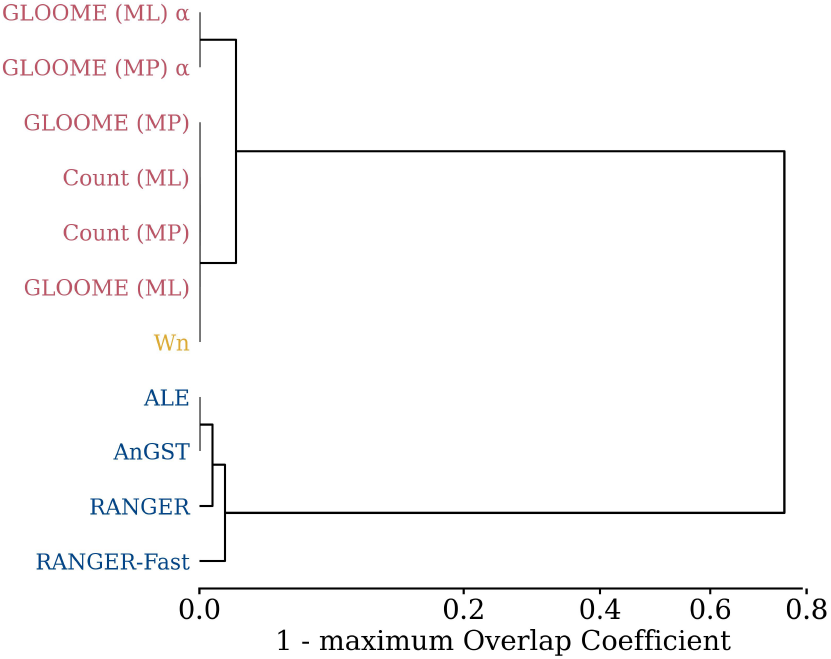
Clustering analysis of the inferred sets of HGT events. Shown here is a UPGMA dendrogram of the tested methods based on their maximum Overlap Coefficient. Overlap Coefficient is the ratio of the size of the intersection to the size of the smaller set (of inferred HGT events) in the pair of methods. Maximum Overlap Coefficient is the maximum of this metric across all stringencies of any pair of methods. This is a similarity measure between 0 and 1, and 1 minus this value is used here as a distance measure. Red: implicit phylogenetic methods. Blue: explicit phylogenetic methods. Yellow: parametric method Wn. To enhance clarity, a power transformation with exponent 0.6 was applied to the x-axis, spreading apart internal nodes that would otherwise appear too close to zero. *α*: GLOOME without species tree. Alt text: Dendrogram showing the clustering of the tested methods based on their maximum Overlap Coefficient. The dendrogram shows two main clusters: one for implicit phylogenetic methods (GLOOME and Count) and one for explicit phylogenetic methods (ALE, RANGER, AnGST). The parametric method Wn clusters with the implicit phylogenetic methods.

Supplementary Fig. S1 confirms the very low overlap between implicit and explicit phylogenetic methods shown in Fig. 1. Within these two groups, we generally see relatively low overlap coefficients at higher stringencies but strong overlaps at low stringencies. This indicates that high-confidence inferences tend to be different even across similar methods, while for low-confidence inferences, one set of inferences tends to be a subset of the other for any pair of methods from the same class (implicit or explicit).

### Implicit phylogenetic methods infer a higher percentage of co-acquisitions as neighbors

When a method infers that two genes from different families were acquired on the same terminal branch of the genome tree, we refer to them as co-acquisitions. If both genes are also inferred to originate from the same donor branch, we call them co-transfers. These terms do not imply that the genes were acquired in a single HGT event. However, some such pairs likely indeed stem from the transfer of a single DNA fragment containing multiple adjacent genes, which would initially remain neighbors in the recipient genome (Dilthey and Lercher, 2015, Pang and Lercher, 2017). While independent transfers could also result in neighboring genes by chance, this is expected to be rare, as quantified below.

We do not know in advance how often true co-acquired gene pairs are the result of single HGT events, and how often they should still be neighbors. However, if one method finds a much higher fraction of neighboring co-acquisitions than another, this is unlikely to be the result of chance – instead, it suggests a lower rate of false-positive inferences by the first method.

For each method, the number of inferred co-acquisitions varied based on the stringency used for each inference method. As we decrease the stringency, the number of co-acquisitions inferred by ALE, Wn, Count, and GLOOME (ML) varies from about 100 to 5 × 10^5^. In contrast, GLOOME (ML, without tree), and RANGER infer much higher numbers of co-acquisitions in general, varying from about 2 × 10^4^ to 10^6^ (Supplementary Fig. S2b). For a fair comparison, the percentage of neighbors should be compared at comparable levels of stringency, i.e., at the same number of inferred co-acquisitions.

The percentage of co-acquisitions in which the co-acquired gene pairs are separated by zero or one intervening gene is notably high, but this percentage declines sharply as the number of intervening genes increases (Fig. S5). For the main text, we define two genes as neighbors if they are separated by at most one other gene. Supplementary Table S1 summarizes the results for other values of the maximum number of intervening genes.

At any given number of co-acquisitions – i.e., any given level of stringency – the implicit phylogenetic methods Count (ML or MP) and GLOOME (ML or MP) almost always infer a higher percentage of neighboring co-acquisitions than the explicit phylogenetic methods or Wn (Fig. 2, Table 1). For example, at 10^3^ co-acquisitions, Count (MP) infers more than 8 times the number of neighboring co-acquisitions inferred by ALE. Count and GLOOME make the exact same inferences with the MP approach. Across stringencies, they outperformed all other methods, with GLOOME (ML) taking second place except at the most stringent thresholds. GLOOME and Count rely solely on the presence and absence of genes across the genome tree.

**Table 1.** Count (MP) infers a higher percentage of neighboring co-acquisitions. Shown here are the mean, standard deviation, and maximum of the percentage of neighboring co-acquisitions, as well as the stringency at which the maximum occurs. This table shows only *t* = 1, where *t* is the maximum number of intervening genes between two co-acquired genes to be considered neighbors. Supplementary Table S1 shows the same for higher values of *t*. Methods using presence-absence outperform the others for all tested values of *t*, with Count (MP) leading among them. Methods such as ALE or Wn have lower variation across their stringencies and low mean percentages of neighboring co-acquisitions. *α* : GLOOME without species tree

| Method | Mean | std | Max | Stringency |
| --- | --- | --- | --- | --- |
| Count (MP) | 1.893 | 1.554 | 4.737 | 7.000 |
| GLOOME (MP) | 1.893 | 1.554 | 4.737 | 7.000 |
| GLOOME (ML) | 1.076 | 0.491 | 1.920 | 0.956 |
| Count (ML) | 0.896 | 0.523 | 2.600 | 1.000 |
| GLOOME (MP) $\alpha$ | 0.896 | 0.409 | 1.376 | 8.000 |
| GLOOME (ML) $\alpha$ | 0.499 | 0.119 | 0.711 | 1.000 |
| Wn | 0.470 | 0.489 | 1.613 | 13.000 |
| ALE | 0.452 | 0.158 | 0.750 | 0.770 |
| AnGST | 0.337 | 0.000 | 0.337 | 1.000 |
| RANGER-Fast | 0.334 | 0.001 | 0.336 | 1.000 |
| RANGER | 0.319 | 0.036 | 0.409 | 1.000 |

**Fig. 2.**
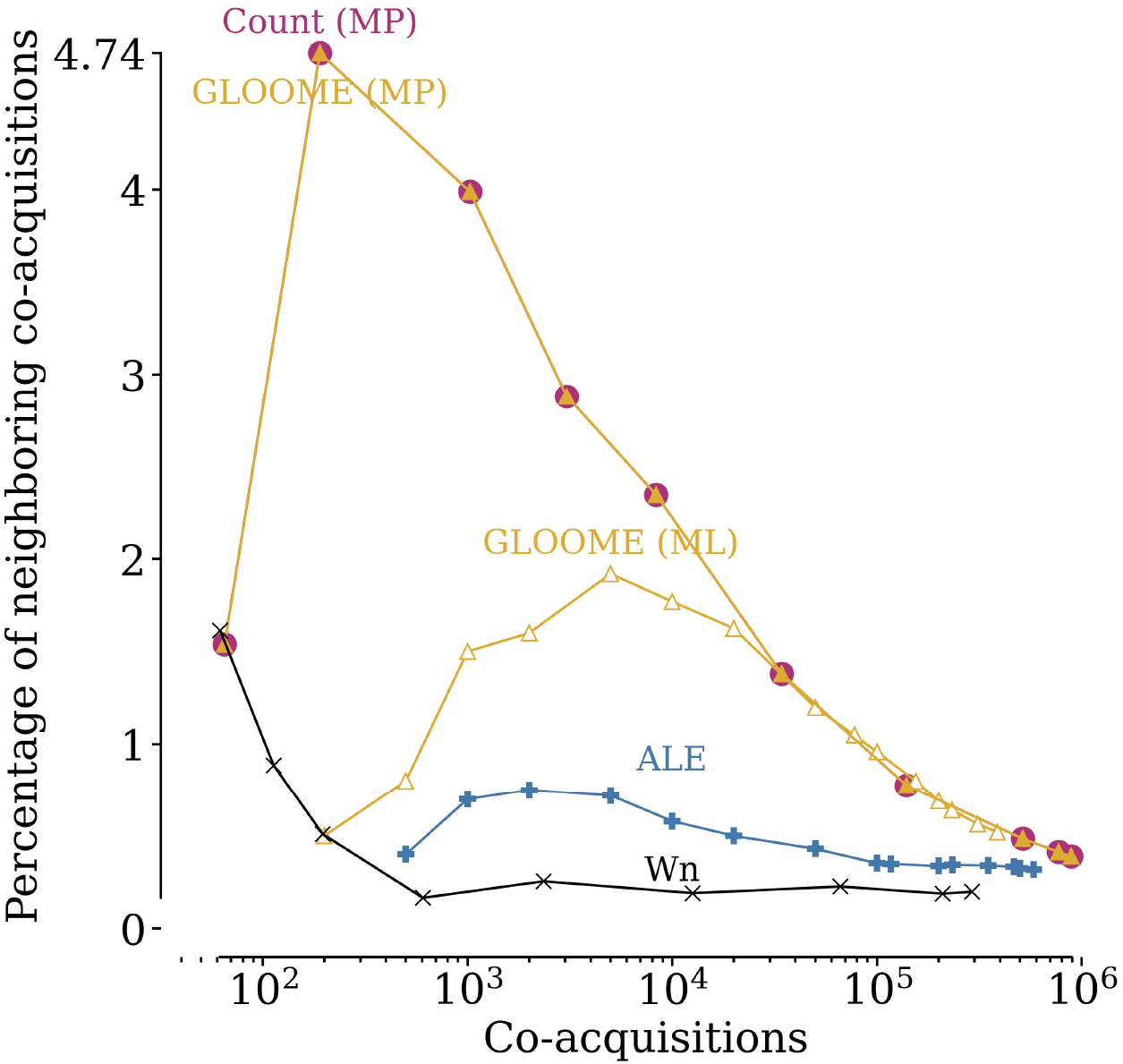
Count (MP) and GLOOME (MP) outperform other methods at inferring neighboring co-acquisitions. The y-axis shows the percentage of neighboring co-acquisitions inferred. Higher stringencies of individual methods are expected to lead to fewer false positive predictions; to make results comparable across methods, the x-axis indicates the number of co-acquisitions. Additional methods are shown in Supplementary Fig. S2b. Alt text: Figure shows line plots of the percentage of neighboring co-acquisitions inferred by several methods on the y-axis, vs the number of co-acquisitions inferred (at different stringency values) on the x-axis.

If one or both genes inferred to have been co-acquired by a given method are false inferences, or if they result from independent HGT events, there is no reason to expect them to be genomic neighbors. Indeed, the expected percentage of neighboring pairs among randomly located genes is very low – typically between 0.09% and 0.18%, with most values around 0.13% (Figure S2a). Correcting for this background expectation does not alter the results.

It appears likely that gene trees are more accurate when they are based on more sequence information, that is, for longer gene sequences. As expected, methods that utilize the gene tree information perform slightly better on longer genes (Supplementary Fig. S3); however, the explicit phylogenetic methods still infer lower percentages of neighboring co-acquisitions compared to the well-performing implicit methods. In sum, the use of gene trees appears to lead to a higher fraction of false-positive HGT inferences.

Gene acquisitions inferred on short terminal branches of the species tree must be (relatively) recent, while gene acquisitions inferred on long terminal branches can be recent, but can also be much older. Thus, we expect that the events inferred for short branches are on average younger than those inferred for long branches. As shown in Supplementary Fig. S4, method performance ranking varies little with branch length, suggesting that our results generalize across phylogenetic depths.

With GLOOME, one can also infer HGTs without providing a *species tree*, in which case the method will infer a tree topology that is most consistent with the phyletic patterns. Without an input species tree, GLOOME consistently performs worse, inferring lower percentages of neighboring co-acquisitions, than the corresponding case of ML or MP with an input species tree (Fig. 3)

**Fig. 3.**
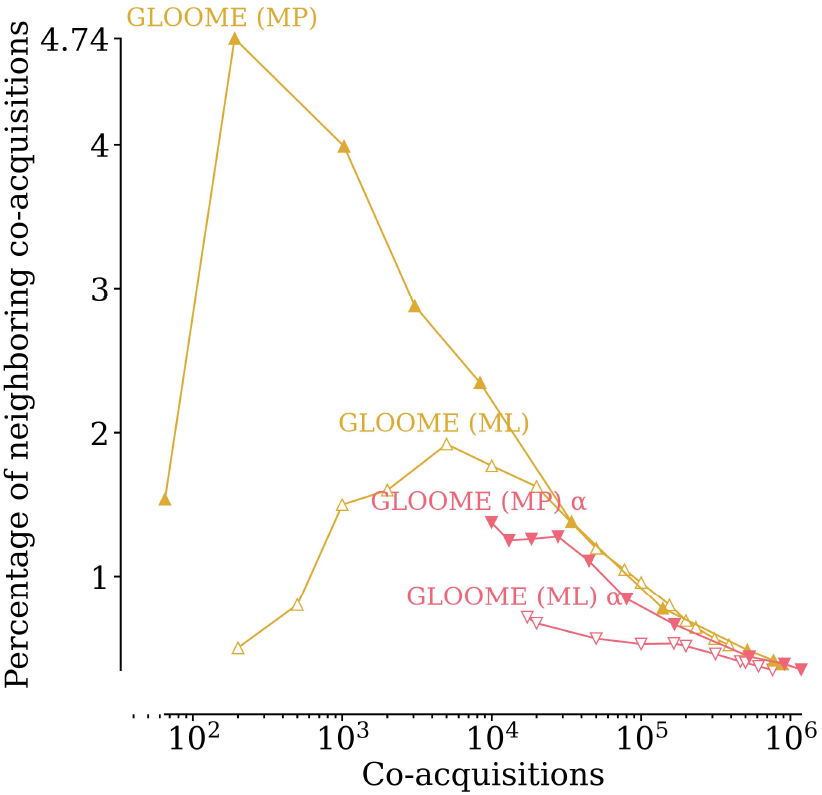
GLOOME performs worse when run without a species tree. Shown here is the percentage of neighboring co-acquisitions inferred by GLOOME (ML) and GLOOME (MP) with and without a species tree (the latter is shown in red and marked with a *α*). Alt text: Figure shows line plots of the percentage of neighboring co-acquisitions inferred by GLOOME (ML, MP, with and without species tree) on the y-axis, vs the number of co-acquisitions inferred (at different stringency values) on the x-axis.

We note here that given the small percentages of neighboring co-acquisitions, one must be cautious in interpreting the results. For example, in Fig. S2b, the observed percentage of neighboring co-acquisitions for several methods (e.g., Count (ML), Wn) increases from 0.5% to 1% when we go from about 200 co-acquisitions to 100 co-acquisitions. This means that the same, single neighboring co-acquisition is inferred by the respective method, but the total number of co-acquisitions inferred is halved. Such effect of small numbers greatly affects the apparent performance of the methods. This type of effect also explains why the parametric method Wn, which generally performs worse than all other methods, appears to improve in performance at very high stringencies (Fig. 2).

The explicit phylogenetic methods in our analysis — ALE, RANGER, and AnGST – also provide information on the donors of the acquired genes. Genes acquired on the same branch of the genome tree could originate from the same donor or from different donors. We have no strong *a priori* expectation about the relative probability of these two types of events. However, it appears unlikely that a method would infer the same donor just by chance. Thus, if we see notable differences between methods in the fraction of inferred co-acquisitions that are also inferred to be co-transfers, then this indicates *a posteriori* that (i) some co-acquisitions are indeed co-transfers and that (ii) the method inferring a higher fraction of co-transfers has a lower false-positive rate. Thus, just like for neighboring co-acquisitions, we also expect that more reliable methods should infer more co-transfers among co-acquisitions; further, they should also infer more neighboring co-transfers among co-acquisitions.

Where results are available for both explicit phylogenetic methods, RANGER almost always infers higher percentages of co-transfers among co-acquired genes than ALE (Fig. 4a). At increasing stringencies for ALE – where no RANGER results are available – ALE infers higher percentages of co-transferred genes.

**Fig. 4.**
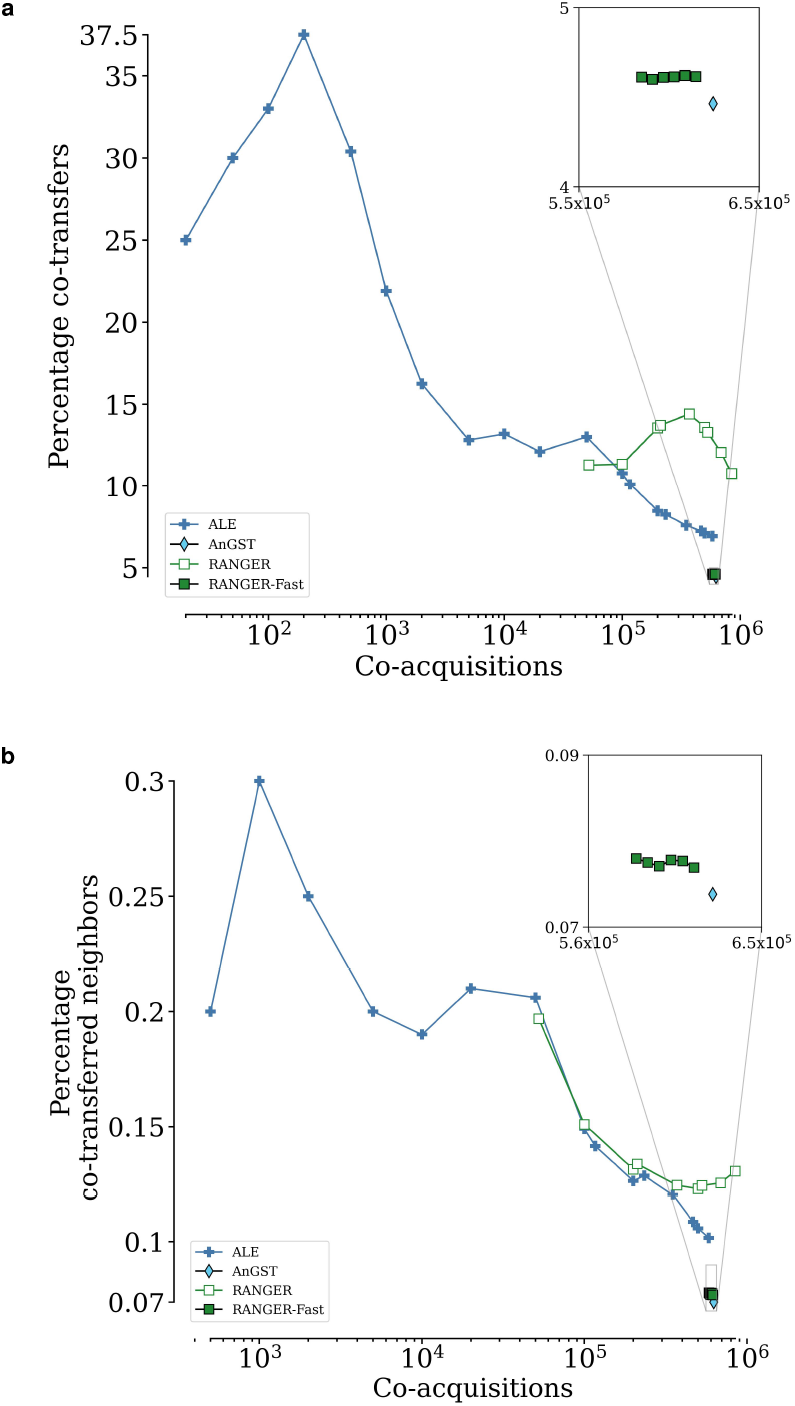
RANGER performs better or similar to ALE at inferring co-transferred genes and co-transferred neighbors. At lower stringency, RANGER infers higher (a) percentages of co-transfers and (b) percentages of co-transferred neighbors than ALE. Although at higher stringency, ALE generally makes better inferences, it is either similar to RANGER or the performance can not be compared as no data from RANGER is available. AnGST and RANGER-Fast generally provide much worse inferences than ALE and RANGER. Note that only explicit phylogenetic methods are shown here, since other methods do not provide information about the source of transfer. Alt text: Two subfigures both showing the number of co-acquisitions on the x-axis and on the y-axis, (a) the percentage of co-transfers and (b) the percentage of co-transferred neighbors, as line plots with one line for each method.

In terms of the inferred percentage of co-transferred neighbors among co-acquisitions, RANGER and ALE perform similarly (Fig. 4b). The results for ALE at higher stringency – where we have no data from RANGER – can again be attributed to the effects of small numbers. Given that the performance of the two methods in this test is similar, one may choose ALE over RANGER simply because it is faster to run. In our analyses, ALE finished running on the dataset using 100 threads in ≈ 18 minutes, whereas RANGER took ≈4.5 hours. AnGST and RANGER-Fast, while faster, perform much worse: they consistently infer very similar percentages of co-transferred genes and co-transferred neighbors, which are substantially lower than those inferred by ALE and RANGER.

At relatively high stringencies (*<* 10^4^ co-acquisitions), ALE infers a relatively large proportion (*>* 13%) of gene pairs as co-transfers (Fig. 4a). According to Fig. 4b, only about 0.2% of co-acquired gene pairs are co-transferred neighbors at this stringency. This indicates that only about 1.5% (i.e., 0.2%/13%) of co-transferred genes are neighbors. This low fraction is roughly consistent with Fig. 2, where about 0.5% of gene pairs inferred by ALE to be co-acquired are neighbors. The low fraction of neighbors indicates that the majority of what we call co-transfers (and, consequently, also co-acquisitions) are the result of independent HGT events. Importantly, that does not invalidate our basic assumption: that a higher fraction of inferred neighboring co-acquisitions (or co-transfers) indicates that the corresponding inference method is more reliable. Instead, it just emphasizes that a perfect inference method would not predict that 100% of co-acquisitions are neighbors.

## Discussion

The present study is the first systematic comparison of phylogenetic methods for inferring HGT. Our analysis reveals several important insights that can guide the choice of method for HGT inference in a given biological study.

Implicit phylogenetic methods, which rely solely on phylogenetic profiles, more accurately capture the spatial clustering of horizontally transferred genes in recipient genomes compared to explicit phylogentic methods. The latter infer HGT events from topological conflicts between gene and species trees, making them vulnerable to gene tree reconstruction errors that can inflate false positive rates for acquisitions and co-acquisitions. The idea that gene tree noise undermines the reliability of explicit methods is supported by the slight performance increase of the explicit phylogenetic methods for long compared to short genes (Supplementary Fig. S3). This aligns with earlier studies showing that statistical errors in gene tree reconstruction can cause topological incongruities with species trees, leading to spurious HGT inferences. Including low-confidence branches exacerbates this problem, increasing false positive rates (Than et al., 2007, 2008). In contrast, implicit methods bypass sequence-level reconstruction and instead rely on gene family assignments, which are considerably less error-prone.

It is surprising to find that the maximum parsimony approaches (of Count and GLOOME) perform on par or often better than their maximum likelihood counterparts, given that maximum likelihood is often considered a superior approach, e.g., in phylogeny reconstruction (Gadagkar and Kumar, 2005). The difference between the two versions of Count is larger when the stringency is higher. The strong increase in the percentage of neighboring co-acquisitions indicates that the strategy of increasing Count (MP)’s stringency by increasing the gain/loss penalty ratio works well, leading to fewer, but more accurate HGTs. In contrast, increasing the stringency of maximum likelihood inferences by selecting events with increasingly higher inferred posterior probabilities appears to be less effective. Future work should explore alternative ways to vary the stringency of maximum likelihood inferences.

Among tree reconciliation methods, RANGER and ALE generally performed better or similar to the others at inferring co-transfers and co-transferred neighbors. Given that ALE is aware of extinct taxa (Szöllosi et al., 2012), it is surprising that it does not perform better than RANGER where results are available for both.

Our methodology is – at least in principle – biased in favor of Wn, the only sequence composition-based method included in our study. Wn is based on oligonucleotide frequencies, which may differ systematically across genomic regions. Accordingly, we may expect Wn to often flag neighboring genes as co-acquisitions, regardless of whether they were indeed the result of HGT. Despite this bias, Wn still performs worse than the phylogenetic methods tested.

Keeping these results in mind, we recommend the use of the implicit phylogenetic methods Count or GLOOME to identify horizontal gene acquisitions. In particular, the MP versions of Count and GLOOME outperform all other methods in identifying neighboring co-acquisitions, especially at higher stringencies (gain/loss penalty ratios 4 to 7; Fig. 2). If inferring the donor of HGT events is a requirement, explicit phylogenetic methods must be used. In this case, both ALE and RANGER are adequate choices, although ALE is computationally more efficient than RANGER. ALE can be made more stringent than RANGER by choosing a high stringency (> 0.86 transfers per reconciliation model). At this level of stringency, its inferences are the most reliable among all tested scenarios for explicit methods, as indicated by the highest percentages of neighboring co-acquisitions or co-transfers (Fig. S2c). On that note, for any method used, we generally recommend using a minimum stringency (see Table 1) and focus on a subset of reliable HGT inferences.

Although Gammaproteobacteria are a well-studied and diverse bacterial clade, these findings might vary for other datasets. Specifically, results could differ when analyzing only recent gene transfers or examining shallower phylogenetic trees, such as those limited to *Escherichia coli* strains. However, this seems unlikely, given that the relative performance ranking of the methods remains mostly consistent across branch lengths S4. Because many users will not spend time optimizing most of the parameters offered by the various programs, we ran each program only with their default settings. We note, however, that these parameters can dramatically change their performance.

Our conclusions rely mainly on the analysis of neighboring co-acquisitions and co-transfers. A limitation of this approach is the lack of an *a priori* expectation for how often gene pairs co-acquired on the same branch of the species tree indeed result from the same HGT event, and how often these are expected to still be neighbors in extant genomes. As a result, our method can provide relative rankings of inference methods, but cannot assess their reliability in absolute terms. Nonetheless, the substantial differences between methods in the fraction of neighboring co-acquisitions suggest that some of these co-acquisitions truly reflect single HGT events. If all methods had similar error rates, we would expect similar fractions of neighboring and co-transferred genes across methods – which is not the case. These differences therefore indicate varying reliability and provide a basis for improving HGT inference methods.

## Methods

### Genomic data selection and cleaning

We used the Gammaproteobacterial dataset (NCBI taxonomic ID: 1236) from the EggNOG database v6 (Hernández-Plaza et al., 2023) for orthologous genes (non-supervised orthologous groups, or *NOGs*). The corresponding nucleotide sequence information (for Wn execution) was downloaded from the NCBI GenBank database (Sayers et al., 2022) using the NCBI Datasets CLI program. The reference species tree downloaded from ASTRAL WoL (Zhu et al., 2019). Specifically, the `astral - branch length – cons - astral.cons.nwk` file was used. In brief, this means that the topology of the tree was inferred using ASTRAL and the branch lengths were re-estimated with maximum likelihood, using the 100 most conserved sites per gene. We retain only the taxa belonging to the maximum overlap of taxa between the set of NOGs, such that their sequence data could be retrieved and that these taxa could be reliably mapped onto the species tree. Furthermore, to ensure that gene families were large enough to infer meaningful HGT events, NOGs containing less than 10 genes or less than 30 taxa were removed.

NOGs were then selected around the average sized NOGs such that they together represented all the 359 taxa in the dataset, resulting in a set of 1286 NOGs. The gene trees were taken from the EggNOG database and rooted using Minimum Ancestor Deviation (Tria et al., 2017).

### Inferring HGT

All the programs were executed separately, using default parameters. For ALE, the *ALEml_undated* program was used, which does not require a dated species tree (Szöllősi et al., 2015). RANGER-DTL v2.0 was executed with and without the ‘fast’ option. AnGST was executed with parameters as suggested in the manual.

The stringency of ALE, RANGER, and RANGER-Fast is defined as the mean number of transfers inferred per reconciliation model. At increasing stringency levels, we exclude HGT events that are inferred with lower frequencies across all of the output the reconciliation models. For Count, the ’Asymmetric Wagner Parsimony’ method was used as the maximum parsimony method. Unlike GLOOME where we also ran without a species tree for both ML and MP settings, Count could be run only with a species tree. For both of these implicit phylogenetic methods, the stringency is defined as the gain/loss penalty ratio given as input. Unlike the explicit phylogenetic methods where we vary the stringency on the output of a single run, with the implicit phylogenetic methods we run the program multiple times, each time with a different gain/loss penalty ratio. These phylogenetic HGT inference methods have publicly available implementations that we used with default settings. In contrast, we implemented the parametric method Wn ourselves, as described below, since there is no publicly available implementation. The parametric method Wn was implemented based on the original publication (Tsirigos and Rigoutsos, 2005), and as suggested, we used an oligonucleotide size of 8 nucleotides. Wn calculates a *typicality score* of how similar the gene sequence is to the genome sequence based on the oligonucleotide frequencies in both. These frequencies were efficiently calculated using KMC (Kokot et al., 2017). In the original paper for Wn, the authors used an automatic threshold detection method. This method rank orders the genes based on their typicality scores, smoothens this curve (requiring a rolling window average), takes a derivative of this curve, and then considers transferred genes as those that are above the point where the derivative becomes *approximately* constant. Although this method is termed *automatic*, it still depends on both the rolling window size used for smoothing and the user’s notion of when exactly does the derivative become approximately constant (i.e. the double derivative is approximately zero). Instead of making the choice of when the double derivative is approximately zero, we used multiples of the *median absolute deviation* (MAD) of the double derivative around zero, which also helps to set the stringencies. These multiples represent the *stringency* parameter that goes from 4 times the MAD to 13 times the MAD. Note that it is generally recommended to use at least 2.5 times the MAD, and so we are already being highly strict even when Wn is run at the lowest stringency. The rolling window size was automatically determined by iteratively finding the longest Savitzky-Golay filter such that the root mean square error of the smoothed curve with respect to the original curve stops decreasing. The code for Wn is available at the code repository linked in the Data Availability section.

### Co-acquisition and co-transfer analysis

To limit the effect of small numbers, chromosomes with less than 1000 genes were excluded from calculations of co-acquisitions, co-transfers, and neighbors. At any given stringency of a method, if the number of co-acquisitions was less than 20, that stringency was excluded from the analysis. The co-acquisitions discussed in this study are those where both genes have known positions in their respective chromosomes. In any gene family, genes from EggNOG that could not be retrieved from GenBank, or pair of genes co-acquired by the same taxon but not in the same chromosome/contig in the GenBank data, were excluded from this analysis.

We calculated the expected percentage of neighboring co-acquisitions under the null model that the two genes were acquired in independent HGT events or that one or both HGT inferences are false. We first calculate the expected number of neighboring co-acquisitions for each chromosome individually, based on the chromosome size and the number of inferred co-acquisitions for that chromosome (see below). The randomly expected number of neighboring co-acquisitions is the sum of these expected numbers across chromosomes. The expected percentage of neighboring co-acquisitions is this expected number, divided by the total number of co-acquisitions. Specifically, if *t* is the maximum number of intervening genes between the two genes in the pair (in our case *t* = 1), and *n* is the number of chromosomes among the inferred co-acquisitions for a method, the expected fraction of neighboring co-acquisitions is

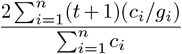

where *c*_*i*_ and *g*_*i*_ are the number of co-acquisitions and the number of genes in chromosome *i*, respectively. This is because for each horizontally transferred gene in a chromosome, the probability of another gene being transferred as a neighbor is 2(*t* + 1)*/g*_*i*_ (i.e. approximating for boundary genes, there are *t* + 1 neighboring positions on either side of a gene out of *g*_*i*_ positions in the chromosome). The expected number of neighboring co-acquisitions is then this probability multiplied by the number of co-acquisitions for that chromosome. The expected *fraction* of neighboring co-acquisitions is the sum of the expected number of neighboring co-acquisitions (per chromosome) divided by the sum of co-acquisitions for all chromosomes.

The expected percentages differ across methods because they are calculated from the number of inferred HGTs for each given genome, considering the genome size. Since these numbers differ across methods, there is some variation across methods in the overall expected number of neighboring co-acquisitions.

## Acknowledgements

We thank Tal Dagan for discussions about the evaluation of HGT inference methods. We thank Carolina Fanalista for her assistance in setting up the project in its early stages.

## Data availability

All of the code used in this study is available at the following Gitlab repository: https://gitlab.cs.uni-duesseldorf.de/general/ccb/hgt_inference_comparative_study The data used in this study can be retrieved from Zenodo: https://doi.org/10.5281/zenodo.14781414.

