## Supplementary Information for "Horizontal Gene Transfer Inference: Gene presence-absence outperforms gene trees"

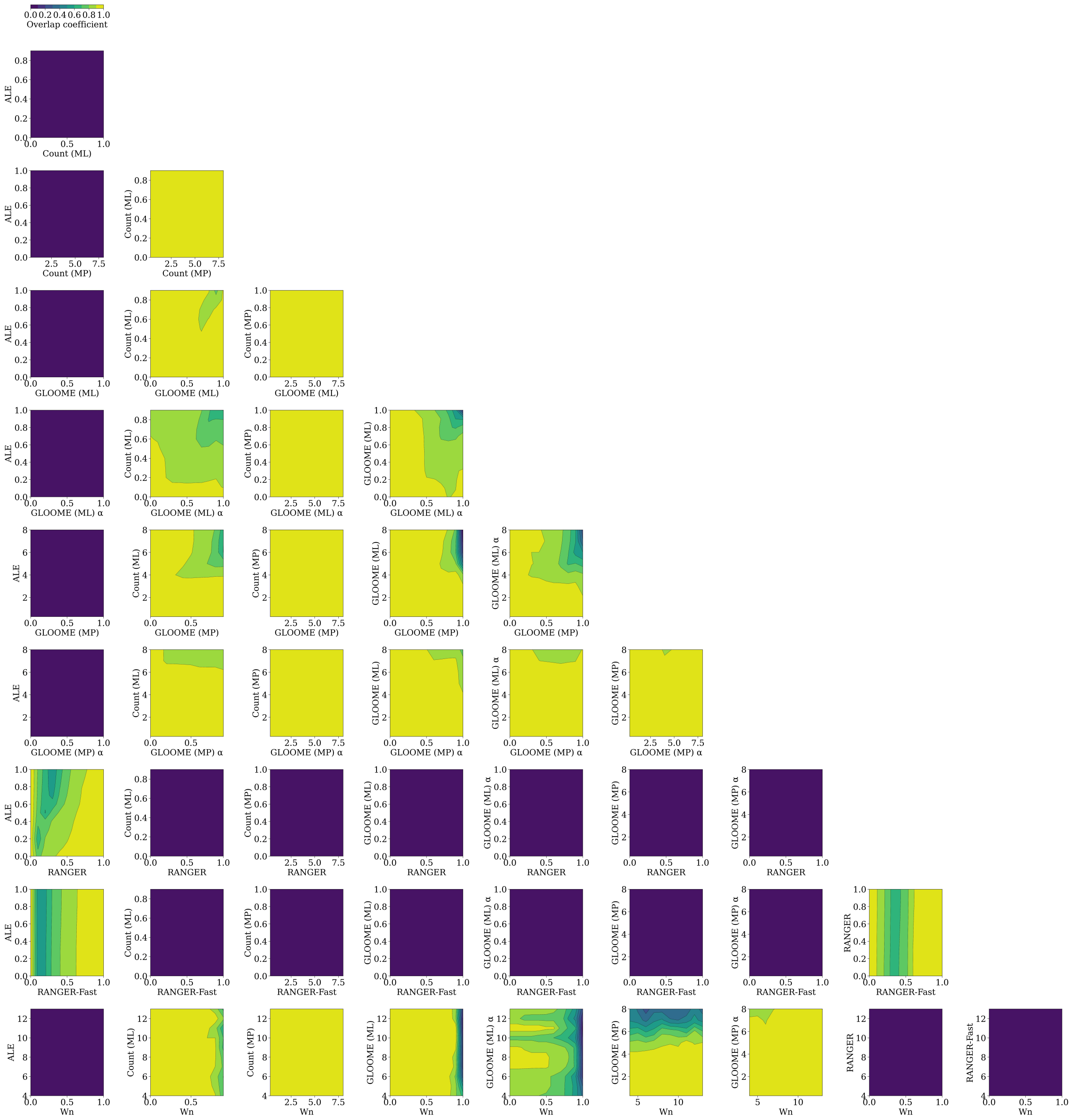


**Fig. S1. Contour plots of Overlap Coefficient between pairs of inference methods.** Figure shows the Overlap Coefficient between pairs of inference methods, for all gene families in our dataset. Each axis subplot is for one pair of methods, and each axis in each subplot shows stringency values for the corresponding method in the axis label. Colors indicate the Overlap Coefficient between the two methods, with warmer colors indicating higher overlap. The Overlap Coefficient is calculated as the size of the intersection of two sets divided by the size of the smaller set. Here, the two sets are sets of inferred HGT events. Implicit and explicit phylogenetic methods have very low overlap, while within these two categories pairs of methods generally have higher overlap at lower stringencies.
Alt text: Figure showing contour plots for each pair of methods. Method names are axis labels and the stringency values are axis ticks. The color of the contour plots indicates the overlap coefficient between the two methods. $\alpha$ : GLOOME without species tree.

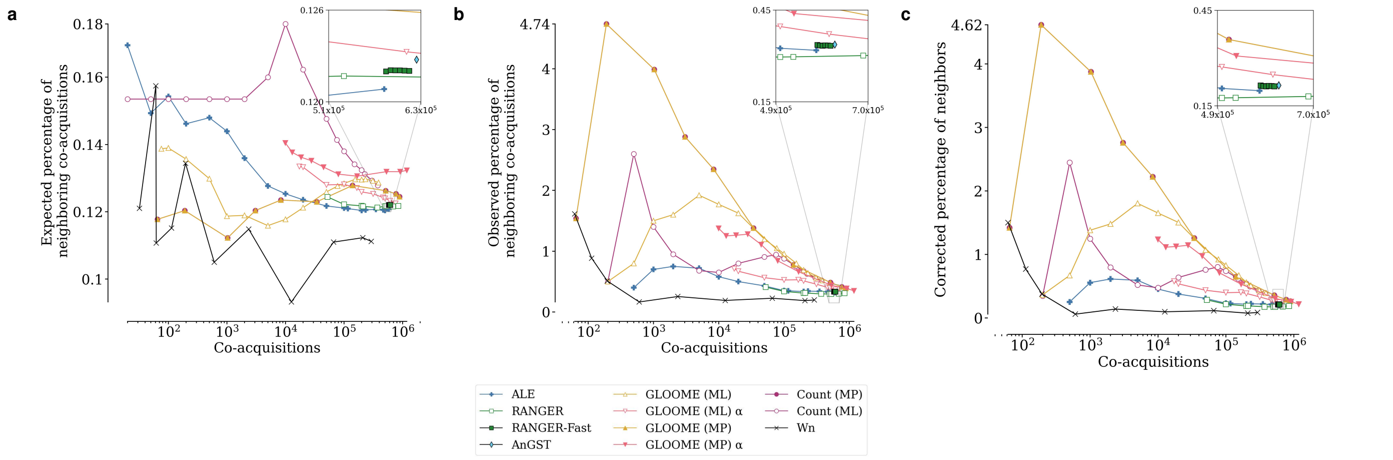


**Fig. S2.** (a) Expected, (b) observed, and (c) observed minus expected percentage of neighboring co-acquisitions, vs number of co-acquisitions inferred. The expected percentage of neighboring co-acquisitions changes with the number of co-acquisitions inferred at various thresholds, and varies between methods, since it depends on the set of chromosomes that those co-acquisitions are inferred on.
Alt text: Figure showing subplots for expected, observed, and observed minus expected percentage of neighboring co-acquisitions respectively. The x-axis shows the number of co-acquisitions inferred, and the y-axis shows the percentage of neighboring co-acquisitions. The legend shows the methods used for inference. $\alpha$ : GLOOME without species tree.

| Method | Mean (t=0) | std (t=0) | Max (t=0) | Mean (t=2) | std (t=2) | Max (t=2) | Mean (t=3) | std (t=3) | Max (t=3) |
|---|---|---|---|---|---|---|---|---|---|
| Count (MP) | 1.324 | 1.187 | 3.684 | 2.198 | 1.666 | 4.737 | 2.396 | 1.716 | 4.961 |
| GLOOME (MP) | 1.324 | 1.187 | 3.684 | 2.198 | 1.666 | 4.737 | 2.396 | 1.716 | 4.961 |
| GLOOME (ML) | 0.657 | 0.335 | 1.180 | 1.347 | 0.593 | 2.380 | 1.547 | 0.692 | 2.840 |
| GLOOME (MP) $\alpha$ | 0.544 | 0.255 | 0.814 | 1.192 | 0.549 | 1.888 | 1.410 | 0.640 | 2.250 |
| Count (ML) | 0.460 | 0.256 | 1.200 | 1.124 | 0.641 | 3.200 | 1.355 | 0.739 | 3.800 |
| Wn | 0.330 | 0.501 | 1.613 | 0.573 | 0.493 | 1.613 | 0.654 | 0.447 | 1.613 |
| ALE | 0.314 | 0.160 | 0.600 | 0.578 | 0.165 | 0.880 | 0.672 | 0.170 | 1.020 |
| GLOOME (ML) $\alpha$ | 0.289 | 0.065 | 0.405 | 0.672 | 0.171 | 0.995 | 0.817 | 0.212 | 1.232 |
| AnGST | 0.196 | 0.000 | 0.196 | 0.454 | 0.000 | 0.454 | 0.561 | 0.000 | 0.561 |
| RANGER-Fast | 0.194 | 0.001 | 0.195 | 0.444 | 0.001 | 0.446 | 0.546 | 0.001 | 0.547 |
| RANGER | 0.186 | 0.026 | 0.250 | 0.425 | 0.046 | 0.539 | 0.528 | 0.059 | 0.673 |

**Table S1.** Mean, standard deviation, and maximum of the percentage of neighboring co-acquisitions inferred by each method, for $t \neq 1$, where $t$ is the maximum number of intervening genes between two co-acquired pair of genes to be considered neighbors.
$\alpha$ : GLOOME without species tree.

| Short gene length | | Long gene length | |
|---|---|---|---|
| Method | Mean | Method | Mean |
| Count (MP) | 1.459 | Count (MP) | 1.556 |
| GLOOME (MP) | 1.459 | GLOOME (MP) | 1.556 |
| GLOOME (ML) | 1.183 | GLOOME (MP) $\alpha$ | 1.128 |
| GLOOME (MP) $\alpha$ | 1.002 | GLOOME (ML) | 1.100 |
| Count (ML) | 0.694 | Count (ML) | 0.900 |
| GLOOME (ML) $\alpha$ | 0.530 | GLOOME (ML) $\alpha$ | 0.646 |
| ALE | 0.324 | ALE | 0.607 |
| RANGER | 0.274 | RANGER | 0.449 |
| Wn | 0.225 | | |

**Table S2.** Ranking of methods according to mean percentages of neighboring co-acquisitions inferred by each method, for gene families with mean gene length shorter or longer than the median value of 799.86 bp. Only co-acquisitions in the range of $10^3$ to $10^5$ were considered, to exclude the effect of small numbers at lower number of co-acquisitions and the effect of low confidence inferences at higher number of co-acquisitions. Note that the relative ranking of the methods remains similar (see also Supplementary Figure S3). $\alpha$ : GLOOME without species tree.

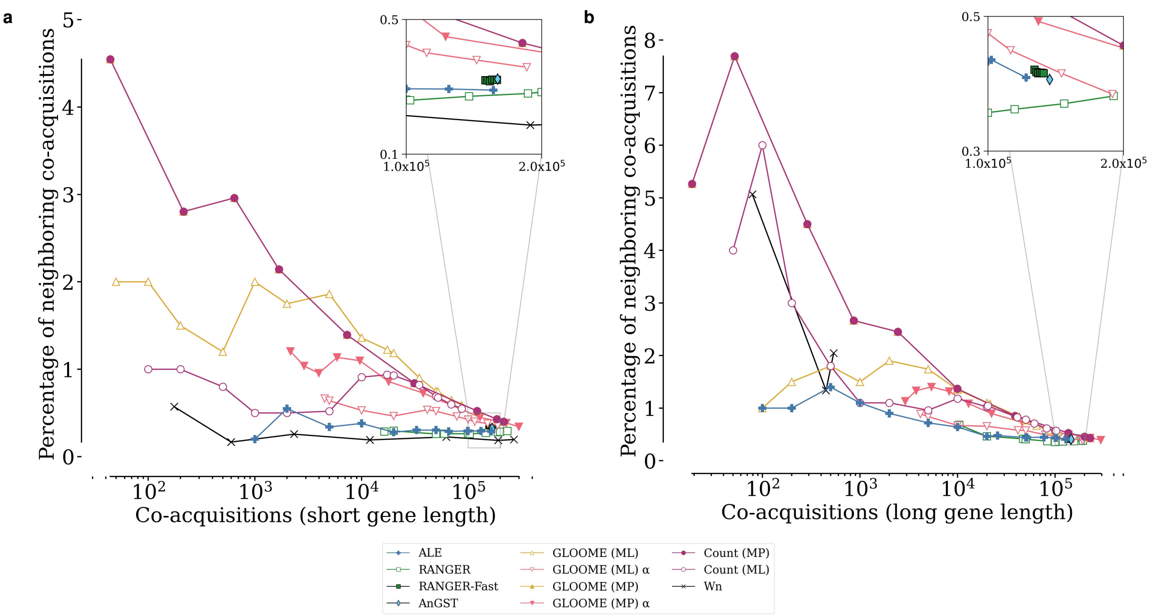


**Fig. S3.** Percentage of neighboring co-acquisitions vs number of co-acquisitions inferred, shown separately for gene families with mean gene length (a) shorter and (b) longer than the median value of 799.86 bp (i.e., the lower and upper halves of the gene length distribution in our dataset). The mean gene length is calculated as the average length of all genes in a gene family. Explicit phylogenetic methods perform slightly better on longer genes. For example, at 2000 co-acquisitions, the percentage of neighboring co-acquisitions inferred by ALE is approximately 0.5% for short genes and approximately 1% for long genes. However, the relative ranking of the methods remains similar (see also Supplementary Table S2) $\alpha$ : GLOOME without species tree.

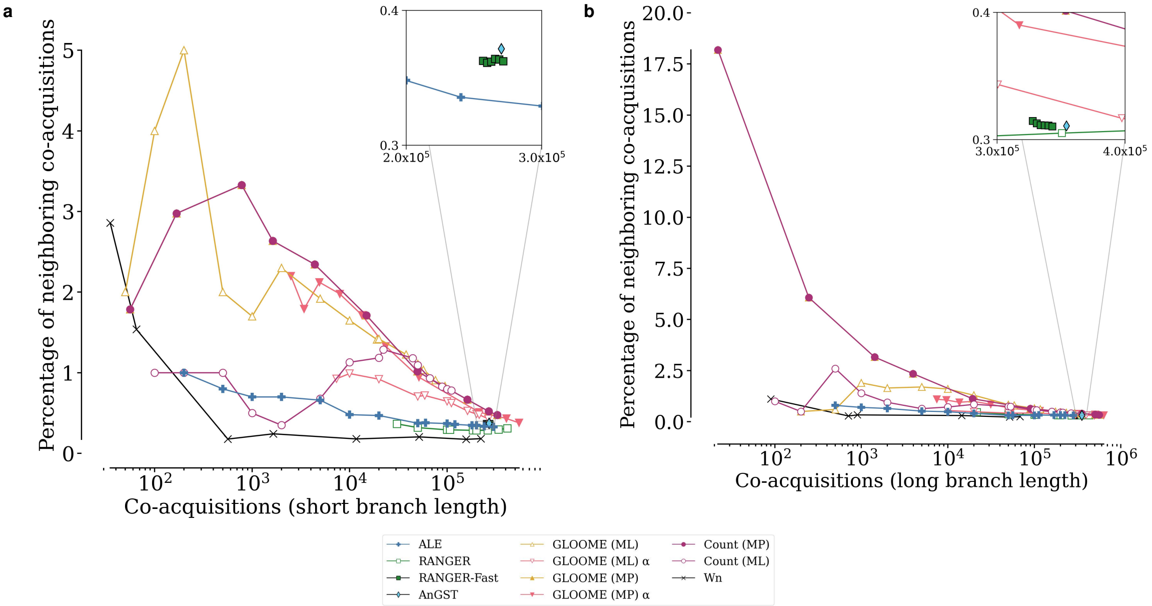


**Fig. S4.** Percentage of neighboring co-acquisitions vs number of co-acquisitions inferred, on species tree branches that are (a) shorter or (b) longer than the median branch length of 0.076 substitutions per site. The relative ranking of the methods remains similar (see also Supplementary Table S3). $\alpha$ : GLOOME without species tree.

| | Short branches | | Long branches |
|---|---|---|---|
| Method | Mean | Method | Mean |
| Count (MP) | 1.927 | Count (MP) | 1.820 |
| GLOOME (MP) | 1.927 | GLOOME (MP) | 1.820 |
| GLOOME (MP) $\alpha$ | 1.726 | GLOOME (ML) | 1.305 |
| GLOOME (ML) | 1.411 | GLOOME (MP) $\alpha$ | 0.933 |
| Count (ML) | 0.906 | Count (ML) | 0.830 |
| GLOOME (ML) $\alpha$ | 0.818 | GLOOME (ML) $\alpha$ | 0.488 |
| ALE | 0.517 | ALE | 0.463 |
| RANGER | 0.326 | RANGER | 0.391 |
| Wn | 0.209 | Wn | 0.259 |

**Table S3.** Ranking of methods according to mean percentages of neighboring co-acquisitions inferred by each method, for species tree branches shorter or longer than the median branch length of 0.076 substitutions per site. Only co-acquisitions in the range of $10^3$ to $10^5$ were considered, to exclude the effect of small numbers at lower number of co-acquisitions and the effect of low confidence inferences at higher number of co-acquisitions. Note that the relative ranking of the methods remains similar (see also Supplementary Figure S4). $\alpha$ : GLOOME without species tree.

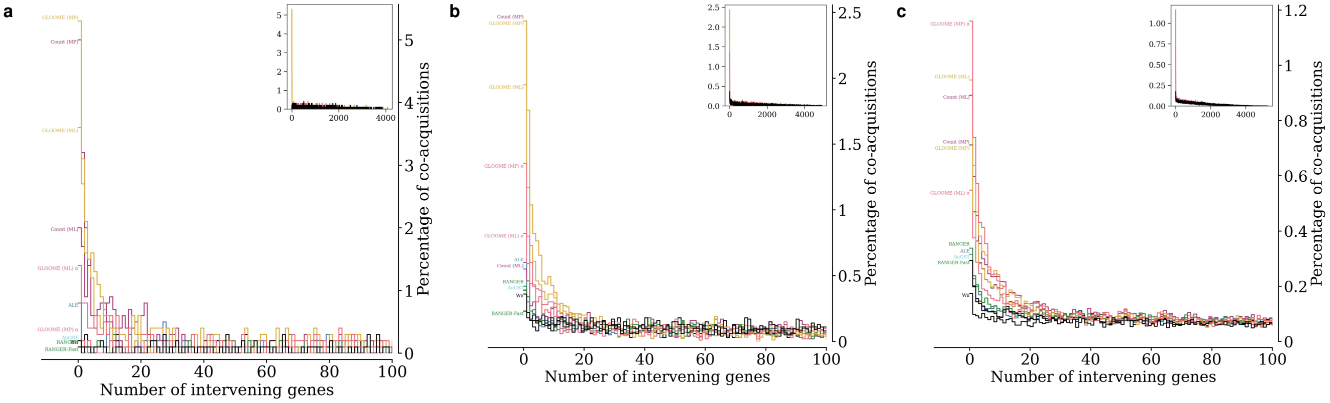


**Fig. S5.** Distribution of distances between co-acquired genes. Shown here are histograms of number of intervening genes between co-acquired genes at three different stringency levels such that the number of co-acquisitions is (a) $10^3$, (b) $10^4$, and (c) $10^5$. The x-axis shows the number of intervening genes between co-acquired genes, and the y-axis shows the corresponding percentage of co-acquisitions. The inset shows the histogram across the full range of intervening genes across all co-acquisitions, while the main plot shows the histogram for the range of intervening genes between 0 and 100. $\alpha$ : GLOOME without species tree.
